# Non-adaptive evolution of the *m. palmaris longus* in the Homininae

**DOI:** 10.1101/015321

**Authors:** Zach Throckmorton, Nicole Forth, Nathanial Thomas

## Abstract

The palmaris longus muscle is widely recognized for its notable variability in living humans. These variations include not only muscle belly reversal, distinct double muscle bellies, duplication and triplication, but also total uni-or bilateral agenesis (absence). A review of the literature and data novel to this study illustrate that different populations of humans exhibit remarkable variation in the frequency of palmaris longus agenesis, from less than 5% of Chinese to nearly 65% of Indians. Comparative dissection-derived data reveal substantial variation in palmaris longus agenesis (PLA) in populations of extant humans (*H. sapiens*), chimpanzees (*Pan spp*.), and gorillas (*Gorilla spp*.) - but not orangutans (*Pongo spp*.), which apparently always develop this muscle. From this pattern, we infer that palmaris longus is undergoing non-adaptive, stochastic evolution in the extant African Homininae, while it continues to have adaptive purpose in *Pongo*, likely due to the orangutans’ greater degree of arboreality than the African apes and humans. Clinical evidence supports this conclusion, at least as it applies to humans. This study highlights the utility of comparative soft tissue data collection and interpretation in elucidating the evolution of anatomical structures that do not fossilize.

## Text

*Musculus palmaris longus*, or simply palmaris longus, is one of the superficial flexor muscles in the antebrachium. It is one of the most variable muscle in the human body, both in its specific form as well as its total agenesis. In humans and in the great apes, this muscle usually arises from the medial epicondyle of the humerus and passes distally, lying medial to the flexor carpi radialis muscle. It continues on to the inferior aspect of the wrist, usually inserting into the flexor retinaculum, and sometimes sending tendinous fibers to the palmar aponerurosis proper. More common variants of palmaris longus include: belly reversal (the muscle belly is located distally rather than proximally), double-belly (both a proximal and a distal muscle belly are present with an intermediate tendon), duplication or triplication (two or three palmaris longus muscles present in one antebrachium), accessory tendinous slips (one palmaris longus muscle that has multiple origins and/or insertions), and agenesis (the palmaris longus muscle is simply absent from the antebrachium, either uni- or bilaterally).

These different variations are potentially clinically relevant as they may cause compartment syndrome (abnormally high pressure in the anterior antebrachial muscle compartment), carpal tunnel syndrome, or Guyon’s syndrome (entrapment of the ulnar nerve in Guyon’s canal, the space between the pisiform and the hamate’s hook). Absence of the palmaris longus muscle has not been shown to significantly affect grip or pinch strength (Sebastin *et al*. 2005, Ertem *et al*. 2007). Its tendon is often harvested for grafts, usually for replacement of the flexor pollicis longus tendon (Unglaub *et al*. 2006). Its use is also documented in the reconstruction of the lower lip in patients with cancer of the lip or gums (Jeng *et al*. 2004), in formation of a sling to correct ptotic (drooping) eyelids (Lam *et al*. 1996), and in the repair of chronic, incomplete tears to the ipsilateral triceps tendon (Scolaro *et al*. 2013). Adverse effects suffered by patients resulting from the surgical removal and relocation of their palmaris longus have not been documented. Therefore, its absence likely does not affect fitness in humans.

In primates, PLA is documented in neither prosimians nor monkeys. However, Hominidae PLA is variable. Because the provenience of dissected apes is rarely recorded in the literature, subpopulation distribution of PLA is understood only in humans (Table 1). Some human populations, such as Chinese, exhibit palmaris longus agenesis frequency of less than 5% (Roohi *et al*. 2007), while others, such as Indians, have palmaris longus agenesis of nearly 65% (Ceyhan & Mavt 1997) – though another study of Indians (Kapoor *et al*. 2008) reported PLA of less than 20%, unsurprisingly indicating that a nation of over 1 billion humans exhibits regional variability. A similar pattern is found in the populous nation of Nigeria (Adejuwon *et al*. 2012, Kayode *et al*. 2008) Interestingly, both South (Machado & Di Dio 1967) and North American (Thompson *et al*. 1921) Natives exhibit low rates of PLA, perhaps owing to their north Asian ancestry. Interestingly, Chilean college students exhibit higher rates of PLA than Native South Americans (Alves *et al*. 2007), perhaps owing to educational access disparities between Natives and more European-descended people.

**Table 1.** Frequencies of palmaris longus agenesis in human subpopulations.

| Reference | N | Bilateral Presence | Unilateral PLA | Bilateral PLA | Ancestry |
| --- | --- | --- | --- | --- | --- |
| Adejuwon *et al.* (2012) | 564 | 414 | 73 | 77 | Nigerian |
| Albay *et al.* (2013) | 58 | 33 | 6 | 19 | Romanian |
| Alves *et al.* (2011) | 200 | 160 | 22 | 40 | Chilean |
| Ceyhan & Mavt (1997) | 7000 | 2688 | 1365 | 2947 | Indian |
| Eric *et al.* (2009) | 800 | 500 | 173 | 127 | Belgium |
| Eric *et al.* (2011) | 542 | 136 | 288 | 118 | Belgium |
| Ertem *et al.* (2007) | 365 | 77 | 206 | 82 | Turkish |
| Gangata (2009) | 890 | 877 | 8 | 5 | Zimbabwean |
| Hussain & Hasan (2012) | 400 | 312 | 67 | 31 | Saudi Arabian |
| Kapoor *et al.* (2008) | 500 | 414 | 46 | 40 | Indian |
| Kayode *et al.* (2008) | 600 | 413 | 75 | 112 | Nigerian |
| Kigera & Mukwaya (2011) | 800 | 766 | 26 | 9 | Ugandan |
| Machado & Di Dio (1967) | 379 | 365 | 4 | 10 | Native Amazonian |
| Mbaka & Ejiwunmi (2009) | 600 | 560 | 34 | 6 | Yoruban |
| Ndou *et al.* (2010) | 201 | 178 | 12 | 11 | Mixed ancestry South African |
| Roohi (2007) | 153 | 144 | 7 | 2 | Chinese |
| Roohi (2007) | 150 | 133 | 12 | 5 | Malay |
| Sebastin *et al.* (2005) | 418 | 377 | 17 | 24 | Chinese |
| Sebastin & Lim (2006) | 329 | 314 | 11 | 4 | Chinese |
| Thompson *et al.* (1921) | 1201 | 908 | 113 | 180 | European-American |
| Thompson *et al.* (1921) | 318 | 300 | 6 | 12 | African-American |
| Thompson *et al.* (1921) | 50 | 48 | 1 | 1 | Native American (Shennecock) |
| Thompson *et al.* (1921) | 101 | 76 | 11 | 14 | Native American (Penobscot) |
| Thompson *et al.* (1921) | 25 | 22 | 3 | 0 | Japanese |
| Throckmorton *et al.* (this study) | 81 | 51 | 16 | 14 | European-American |

There are no systematic, large-sample surveys of muscular variation in the great apes. Reported great ape PLA frequency data are given in Table 2. Though only 13 orangutans are noted for this character in the literature, palmaris longus was present in all 26 of their arms. Of the 61 chimpanzees surveyed and reported by researchers, six exhibited bilateral palmaris longus agenesis and seven exhibited unilateral PLA; that is, approximately one out five of chimpanzees lack palmaris longus in at least one arm. Gorillas exhibit the highest rates of great ape PLA, with 15 of 21 specimens lacking this muscle in one or both antebrachia.

**Table 2.** Frequencies of palmaris longus agenesis in great apes.

| Reference | N | Bilateral Presence | Unilateral PLA | Bilateral PLA |
| --- | --- | --- | --- | --- |
| ***Pongo*** |  |  |  |  |
| Beddard 1893 | 1 | 1 |  |  |
| Chapman 1880 | 1 | 1 |  |  |
| Church 1861-1862 | 1 | 1 |  |  |
| Fick 1895a,b | 1 | 1 |  |  |
| Hepburn 1892 | 1 | 1 |  |  |
| Kallner 1956 | 2 | 2 |  |  |
| Michaelis 1903 | 1 | 1 |  |  |
| Oishi et al 2008,2009 | 3 | 3 |  |  |
| Payne 2001 | 3 | 3 |  |  |
| Primrose 1899,1900 | 1 | 1 |  |  |
| Sonntag 1924a | 1 | 1 |  |  |
| **Totals:** | 16 | 16 |  |  |
| ***Gorilla*** |  |  |  |  |
| Bischoff 1880 | 1 |  |  | 1 |
| Chapman 1878 | 1 |  |  | 1 |
| Deniker 1885 | 1 |  |  | 1 |
| Diogo et al 2011 | 1 |  |  | 1 |
| Duckworth 1904 | 1 |  |  | 1 |
| Eisler 1890 | 1 |  |  | 1 |
| Hartmann 1886 | 1 |  |  | 1 |
| Hepburn 1892 | 2 | 1 |  | 1 |
| Hofer 1892 | 1 |  |  | 1 |
| Macalister 1873 | 1 | 1 |  |  |
| Owen 1868 | 1 | 1 |  |  |
| Pira 1913 | 1 |  |  | 1 |
| Preuschoft 1965 | 3 |  | 1 | 2 |
| Raven 1950 | 1 | 1 |  |  |
| Sarmiento 1994 | 2 | 2 |  |  |
| Sommer 1907 | 1 |  |  | 1 |
| Symington 1889 | 1 |  |  | 1 |
| **Totals:** | 21 | 6 | 1 | 12 |
| ***Pan*** |  |  |  |  |
| Beddard 1893 | 1 | 1 |  |  |
| Carlson 2006 | 2 | 1 | 1 |  |
| Champman 1879 | 1 | 1 |  |  |
| Champneys 1972 | 1 | 1 |  |  |
| Dwight 1895 | 1 | 1 |  |  |
| Gibbs 1999 | 14 | 8 | 3 | 3 |
| Gratiolet & Alix 1866 | 1 | 1 |  |  |
| Hepburn 1892 | 1 | 1 |  |  |
| Humphry 1867 | 2 | 1 |  |  |
| Keith 1899 | 6 | 4 | 1 | 1 |
| Loth 1931 | 10 | 9 | 1 |  |
| MacDowell 1910 | 1 | 1 |  |  |
| Miller 1952 | 1 | 1 |  |  |
| Ribbing 1912 | 1 | 1 |  |  |
| Sarmiento 1994 | 9 | 7 | 1 | 1 |
| Sonntag 1923 | 1 | 1 |  |  |
| Sonntag 1924 | 3 | 2 |  | 1 |
| Swindler & Wood 1973 | 1 | 1 |  |  |
| Tyson 1699 | 1 | 1 |  |  |
| Vrolik 1841 | 1 | 1 |  |  |
| Wilder 1862 | 1 | 1 |  |  |
| Ziegler 1964 | 1 | 1 |  |  |
| **Totals:** | 61 | 47 | 7 | 6 |

Chimpanzees, gorillas, and humans are less arboreal than orangutans (Tuttle and Watts 1985, Hunt 1990, Remis 1995), with the orangutan retaining the most arboreally-adapted musculoskeletal system of the Hominidae. Reliance upon arboreal quadrumanous locomotion necessitates maintenance of strong wrist flexion facilitated by *m. palmaris longus* in orangutans, which have the relatively longest arms of the Homininae and use their hands in a marked variety of postures (Cant 1987). This selection pressure is diminished in more terrestrial apes and humans; that is, negative selection against individuals lacking palmaris longus is relaxed in the African apes and humans. Released from its role in arboreal locomotion, *m. palmaris longus* could potentially be undergoing non-adaptive evolution (*i.e*., drift) in the Homininae. Though our data are not a direct test of this assertion, they are consistent with it. The absence of adverse consequences of surgical removal of palmaris longus in humans suggests the muscle’s vestigiliaty, at least in our own species. *In vivo* electromyographic studies of palmaris longus recruitment in the apes would shed additional light on this issue; the expectation is that the muscle is utilized during typical orangutan arboreal locomotion, but is not important in chimpanzee and gorilla knuckle-walking (and perhaps not recruited when the African apes engage in arboreal behaviors).

While it is tempting to infer that incidence of PLA is a simple linear relationship with each species of Homininae’s degree of arboreality, with the most arboreal orangutans retaining the muscle, the most terrestrial gorilla usually lacking it, and the chimpanzee intermediate for both arboreality and incidence of palmaris longus agenesis, the wide range of human subpopulation PLA frequency variation casts doubt on such a simple interpretation of the data. Some human subpopulations exhibit palmaris longus agenesis frequency like chimpanzees, while others more closely match gorillas. Furthermore, the low sample sizes of the great apes compared to the large sample sizes from many human subpopulations cautions against great confidence in interpreting the evolutionary history of *m. palmaris longus* in the Homininae. Indeed, regarding soft tissue structures that do not fossilize, large-scale surveys of comparative data provide a useful means for understanding anatomical evolution given the currently insufficient understanding of the genetic and developmental etiologies of most specific anatomical structures.

## Acknowledgements

We thank the Lincoln Memorial University DeBusk College of Osteopathic Medicine and Anatomical Sciences Master of Science Program for their support of this research. We also thank the donors who generously gave their bodies to anatomical training and study.

